# Molecular mechanisms of liposome interactions with bacterial envelopes

**DOI:** 10.1101/2023.10.07.561336

**Authors:** Anna Scheeder, Marius Brockhoff, Edward N. Ward, Gabriele S. Kaminski Schierle, Ioanna Mela, Clemens F. Kaminski

## Abstract

Although fusogenic liposomes offer a promising approach for the delivery of antibiotic payloads across the cell envelope of Gram-negative bacteria, there is still limited understanding of the individual nanocarrier interactions with the bacterial target. Using super-resolution microscopy, we characterize the interaction dynamics of positively charged fusogenic liposomes with Gram-negative (*Escherichia coli*) and Gram-positive (*Bacillus subtilis*) bacteria. The liposomes merge with the outer membrane (OM) of Gram-negative bacteria, while attachment or lipid internalization is observed in Gram-positive cells. Employing total internal reflection fluorescence microscopy, we demonstrated liposome fusion with model supported lipid bilayers. For whole *E. coli* cells, however, we observed heterogeneous membrane integrations, primarily involving liposome attachment and hemifusion events. With increasing lipopolysaccharide length the likelihood of full-fusion events was reduced. The integration of artificial lipids into the OM of Gram-negative cells led to membrane destabilization, resulting in decreased bacterial vitality, membrane detachment, and improved co-delivery of Vancomycin—an effective antibiotic against Gram-positive cells. These findings provide significant insights into the interactions of individual nanocarriers with bacterial envelopes at the single-cell level, uncovering effects that would be missed in bulk measurements. This highlights the importance of conducting single-particle and single-cell investigations to assess the performance of next-generation drug delivery platforms.

## INTRODUCTION

Liposomes are highly customizable nano-particles that enhance the therapeutic index of their payload.^1–3^ During the past few years, these nanocarriers have attracted significant attention due to their remarkable potential in global medical healthcare, as evidenced by their crucial role in the COVID-19 pandemic.^4^ However, the clinical translation of liposomal drug delivery systems remains slow, which could be attributed to a limited understanding of the heterogeneous interactions with targeted cells and translational gaps between *in vitro, in vivo* and patient testing.^1,5^

Liposomes are particularly attractive due to their easily modifiable lipid composition, which can achieve various functionalities including membrane fusogenicity, achieved by the inclusion of fusogenic lipids.^6^ The ability to facilitate content delivery across membrane barriers is widely used in eukaryotic cell transfection^6^ and offers great potential in addressing the intrinsic antimicrobial resistance of Gram-negative bacteria.^7–10^ The outer membrane (OM) of Gram-negative bacteria acts as a protective permeability barrier equipped with efflux pumps that prevent many intracellular-acting antibiotics from reaching their target at sufficient concentration.^11,12^ However, through the utilization of antibiotic-loaded fusogenic liposomes (FLs), the active compound is released directly into the periplasmic space. By circumventing the OM, this approach enhances the efficiency of reaching the ultimate target.

To initiate membrane fusion, the liposomes must first attach and engage in close interaction with the encountered membrane. Then, aided by the presence of negative curvature lipids such as 1,2-dioleoyl-sn-glycero-3-phosphoethanolamine (DOPE), lipid mixing between the outer layers is facilitated. This process leads to the formation of stalks and hemifusion intermediates (Fig. 1a). Upon collapse of these intermediates, a fusion pore is formed, resulting in the compartment mixing and the delivery of contents (Fig. 1a). While this pathway has been proven to enhance the antimicrobial activity of various antibiotics,^7–10^ it is essential to highlight that the nanocarrier itself has been reported to affect bacterial cell viability.^13,14^

Currently, the field primarily focuses on enhancing lipid compositions and introducing additional modifications to improve liposomal circulation times,^15,16^ stimuli-responsiveness,^17,18^ specific targeting of bacteria or mammalian cells,^19,20^ and endosomal escape of the delivered cargo,^21,22^ among others. However, the interactions and effects of the vehicle itself on their targets, be it mammalian cells or bacterial envelopes, have not been thoroughly investigated. For instance, while it has been demonstrated that FLs mix their lipids with the OM of Gram-negative bacteria (Fig. 1b), the effects of integrating artificial lipids into the membrane system remain unexplored. Similarly, little is known about the nature of interactions of FLs with systems that are not membrane-enveloped, like the cell wall of Gram-positive bacteria. Despite the compositional mismatch between the lipid-based carrier and the sugar-peptide structure of the envelope, FLs were explored as antimicrobial delivery systems. Although FLs have been reported to interact with the cell wall of Gram-positive bacteria (Fig. 1b), direct evidence for liposomes fusing with the bacterial envelope^7^ or lipid internalization^23^ is still missing.

**Figure 1.**
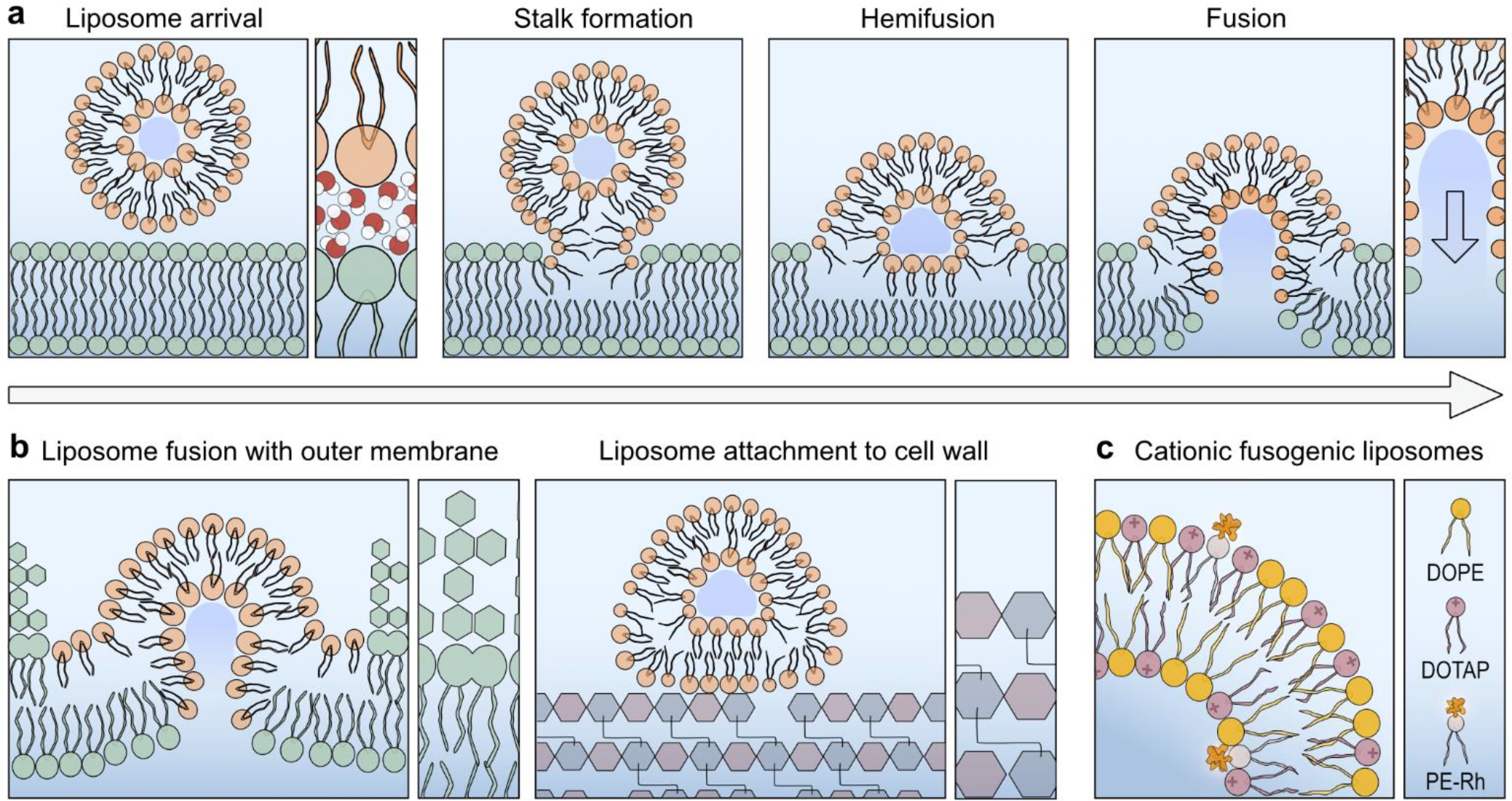
Fusogenic liposomes merge with encountered membranes. (a) Fusogenic liposomes meeting a lipid bilayer must undergo headgroup dehydration to facilitate liposomal attachment and close membrane interactions necessary for membrane fusion. Lipids with high negative curvature, such as DOPE, encourage lipid mixing between the outer monolayers, leading to the formation of a fusion stalk and hemifusion intermediates. In the hemifusion state, the inner monolayers of liposomes and the encountered membrane remain separate. The collapse of this hemifusion state ultimately leads to the creation of a fusion pore and compartment mixing. (b) Schematic depiction of fusogenic liposomes interacting with the outer membrane of Gram-negative bacteria (left) and the peptidoglycan network of Gram-positive bacteria (right). While Gram-negative bacteria are enveloped by an asymmetric membrane of external lipopolysaccharides and inner phospholipids, Gram-positive bacteria are presenting the thick cell wall made from glycan strands and peptides. Fusogenic liposomes merge with the membrane of Gram-negative bacteria while the interactions with the peptidoglycan network of Gram-positive envelopes remain largely unexplored (right). (c) The cFLs composition used in this study comprised the structural lipid DOPE, the cationic co-lipid DOTAP, and the fluorescently labeled lipid Rh-PE. DOPE promotes the formation of liposome-membrane fusion intermediates. DOTAP equips the liposomes with a positive charge that promotes bacterial targeting and facilitates close membrane interactions due to charge attraction. Rh-PE is incorporated to visualize the interactions of cFLs with bacteria using fluorescence microscopy techniques.

To study liposome fusion, bacterial population measurements using lipid-mixing assays are commonly performed on Gram-positive and Gram-negative bacteria. A measurement involves the dequenching and increased fluorescence emission of a reporter embedded within the liposomal membrane when diluted into the target membrane.^7,24^ To complement these population-level measurements, researchers have employed single cell imaging techniques such as epifluorescence microscopy and Transmission Electron Microscopy (TEM).^10,24^ However, while these techniques provide valuable insights, epifluorescence microscopy lacks the resolution to distinguish between fused and attached liposomes and TEM suffers from poor contrast in imaging lipid membranes. Thus, new imaging approaches with higher spatial and temporal resolution are needed to further improve our understanding of the interactions between liposomes and bacterial envelopes in biologically relevant environments. For instance, total internal reflection fluorescence microscopy (TIRF-M) can identify the interaction dynamics of individual nanoparticles with bacterial envelopes,^23^ providing crucial information about the varying susceptibility of individual cells towards FLs.^25^

In this study we make use of a combination of imaging techniques with high temporal and spatial resolution, namely TIRF-M, structured illumination microscopy (SIM), and atomic force microscopy (AFM), to investigate the unknown details of the interaction of cationic fusogenic liposomes (cFLs) with Gram-negative and Gram-positive bacteria. We use super-resolution microscopy to reveal precisely how cFL target Gram-positive *Bacillus subtilis* and Gram-negative *Escherichia coli* bacteria with time-sequenced imaging at the single cell level. cFL were seen to fuse with the OM of Gram-negative cells, while liposome attachment and lipid internalisation was recorded for Gram-positive *B. subtilis* cells. To distinguish between heterogeneous liposome fusion states in real-time, TIRF-M was employed on Gram-negative model bacteria, demonstrating hemifusion and full fusion of liposomes upon bacterial OM encounter. These varying bacterial interactions might affect the efficacy of antibiotic delivery. To understand the effects of lipid addition on cell membrane integrity, we performed cell vitality assays and observed membrane deformations through time-lapse microscopy. The integration of artificial lipids into the OM of Gram-negative cells caused membrane destabilisation and reduced cell vitality. Additionally, we co-delivered Vancomycin, a large antimicrobial agent known for its limited activity against Gram-negative bacteria due to insufficient membrane permeability.^26^ The destabilization of the OM by liposomes enhanced the membrane permeability of free Vancomycin. Besides compromising bacterial integrity, the cationic lipid nanoparticles also caused bacterial aggregation, which, in the context of a blood infection, could lead to severe side effects. With this study, we aim to build a fundamental understanding of nanocarrier interactions with bacterial envelopes to influence the development of next-generation drug delivery systems.

## RESULTS AND DISCUSSION

### Synthesis and characterization of cFLs

Small fusogenic liposomes were synthesized from a 1:1 mole ratio of a structural and a cationic lipid using the well-established thin film hydration technique (Fig. 1c).^27^ The structural, conically-shaped lipid DOPE facilitates lipid mixing between the outer monolayers of liposome and the encountered membrane. The negative curvature of the lipid promotes the formation of a hemifusion intermediate that eventually collapses through the formation of a fusion pore (Fig. 1a)^6,28,29^. Additionally, the cationic co-lipid 1,2-dioleoyl-3-trimethylammonium-propane (DOTAP) was included to facilitate liposome formation and bacterial targeting through electrostatic attractions. Using dynamic light scattering (DLS), the hydrodynamic diameter and the zeta potential of the nanoparticles were measured. The liposomes had a mean hydrodynamic diameter of 184.15 nm (Fig. 2a) and a cationic zeta potential of 23.86 mV ± 1.72 (Fig. 2b). In contrast, the zeta potential of Gram-negative *E. coli* BL21 and Gram-positive *B. subtilis* cells was -15.03 mV ± 1.18 mV and -15.4 mV ± 1.58 mV, respectively (Fig. 2b). This anionic character is a result of phosphate groups in the OM and the phosphodiester bonds within the cell wall.^30^ The coulomb interactions between cationic lipids and the anionic bacterial envelope facilitates membrane encounter and lipid headgroup dehydration, promoting the close membrane interactions required for liposome fusion.^28^

**Figure 2.**
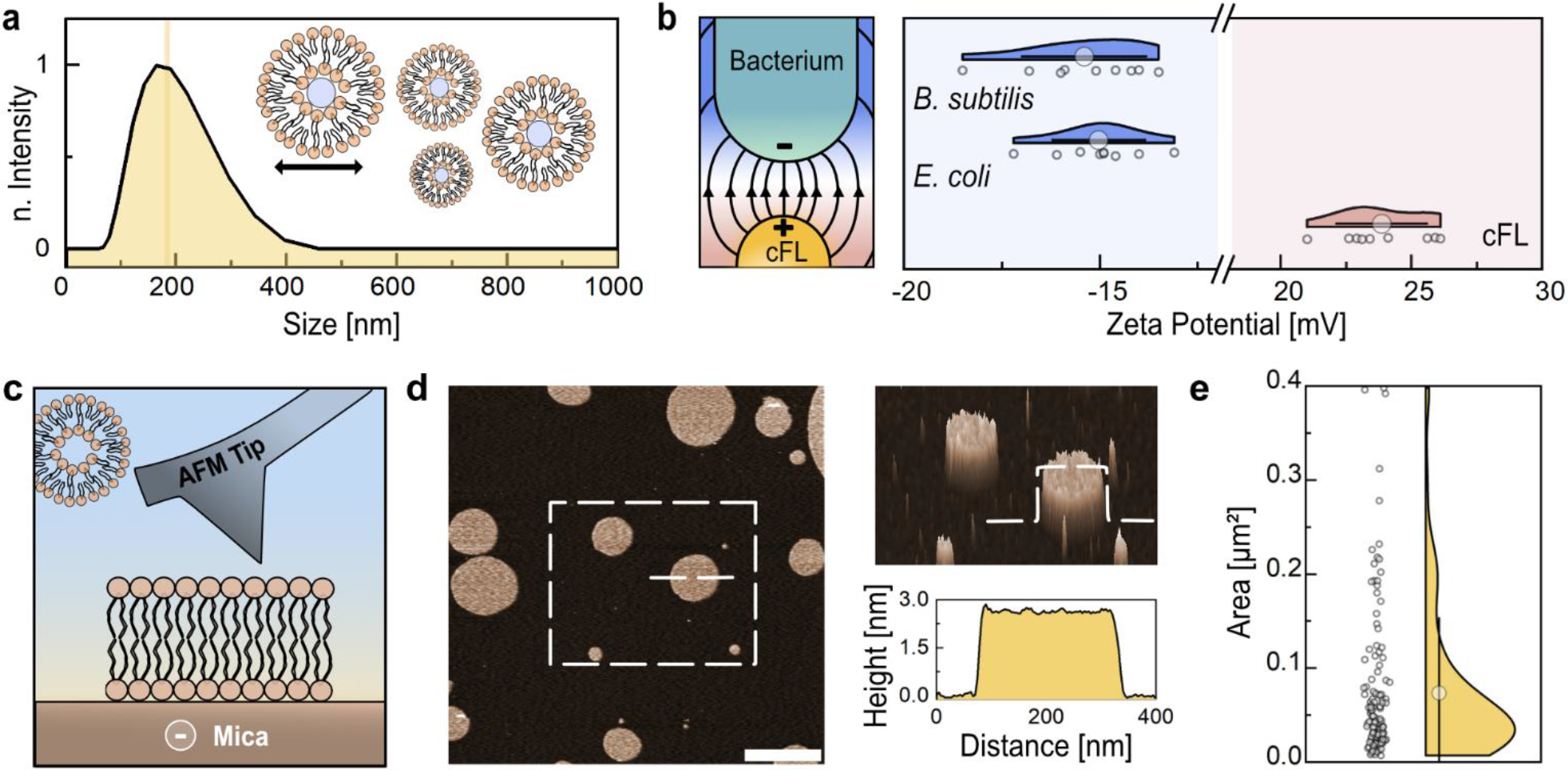
Cationic fusogenic liposomes for charge-driven interactions with negatively charged surfaces. (a) cFLs have a mean hydrodynamic diameter of 184.15 nm. The hydrodynamic diameter was measured using dynamic light scattering. (b) The Zeta potential in mV of cFLs (23.86 mV ± 1.72 mV, *n* = 3), Gram-negative *E. coli BL21* DE3 (BL21) (−15.03 mV ± 1.18 mV, *n* = 3) and Gram-positive *B. subtilis* cells (−15.4 mV ± 1.58 mV, *n* = 3) was measured in 1x PBS. The Schematic illustration on the left displays the coulomb interaction between the cationic FLs and the anionic bacterial envelope that drive bacterial targeting. (c) Schematic depiction of atomic force microscopy on lipid bilayer patches on a negatively charged mica substrate. Representative AFM topography image of cFLs spreading on a mica sheet with bilayer patches of roughly 2.6 nm height. Scale bar, 400 nm. (e) Surface area of cFLs patches. Displaying Mean ± SD, *n* = 129.

We first studied the interactions of cFLs with highly negatively charged surfaces, such as the silicate mineral mica (Fig. 2c). After exposing the material to a nanomolar concentration of liposomes, AFM images were acquired that revealed spherical lipid bilayer patches of ∼2.6 nm height (Fig. 2d). This height profile is consistent with full liposome rupture and formation of a DOTAP:DOPE bilayer.^31^ The bilayers extended over a mean area of 0.073 µm^2^ (95% CI [0.059, 0.087]) (Fig. 2e), which is two orders of magnitude smaller than the average surface area of a bacterium (approximately 6 µm^2^)^32^. Thus, individual liposomes rupture on negatively charged surfaces but they provide insufficient material to fully envelope the surface of a bacterium. However, in contrast to the rigid cell wall of Gram-positive cells, the Gram-negative cell surface consists of an OM. When the liposomes encounter the OM, they might get incorporated, resulting in the mixing of lipid contents of both fluid membrane compartments. To study the membrane fusogenicity of cFLs and the resulting lipid dilution within the host cell membrane, TIRF-M experiments were conducted with *E. coli* total lipid extract supported lipid bilayers (SLB). The formation of the SLB on glass was confirmed with AFM (Supplementary Fig. 1). These membranes were exposed to cFLs that were fluorescently labeled through the incorpora-tion of the lipid 1,2-dioleoyl-sn-glycero-3-phosphoethanolamine-N-(lissamine rhodamine B sulfonyl) (Rh-PE). The imaging plane was focused on the SLB, and upon the arrival of liposomes, a two-stage interaction with the membrane was observed. First, liposomes attached, indicated by the appearance of high-intensity fluorescent areas, followed by radial diffusion of the fluorescently labeled lipid Rh-PE into the bacterial SLB (Fig. 3a, Supplementary Video 1). The measured mean lipid diffusion area was 40.1 µm^2^ (95% CI [26.3, 53.9]) which is approximately six times bigger than the surface area of the bacterium (approximately 6 µm^2^)^32^ and therefore theoretically sufficient to affect the entire OM of Gram-negative cells (Fig. 2b). With lipid diffusion, the fluorescence intensity at the attachment site decayed below 25 % of the initial fluorescence intensity (Fig. 3a,c). During the hemifusion intermediate, the inner monolayers of liposomes and SLB remain separated (Fig. 1a), leaving approximately 50 % of the fluorescently labeled lipids in the inner monolayer behind.^33^ Since the mean residual fluorescence intensity of the cFLs after membrane encounter was less than 50% (Fig. 3c), the cFLs fully merged and mixed their lipids with the targeted SLB. These observations agreed with previous TIRF-M imaging on virus fusion^34^ and cubosome^35^ fusion with SLBs. Control experiments performed on bare glass slides showed liposome attachment and spreading with negligible photo-bleaching effects (Supplementary Fig. 2).

**Figure 3.**
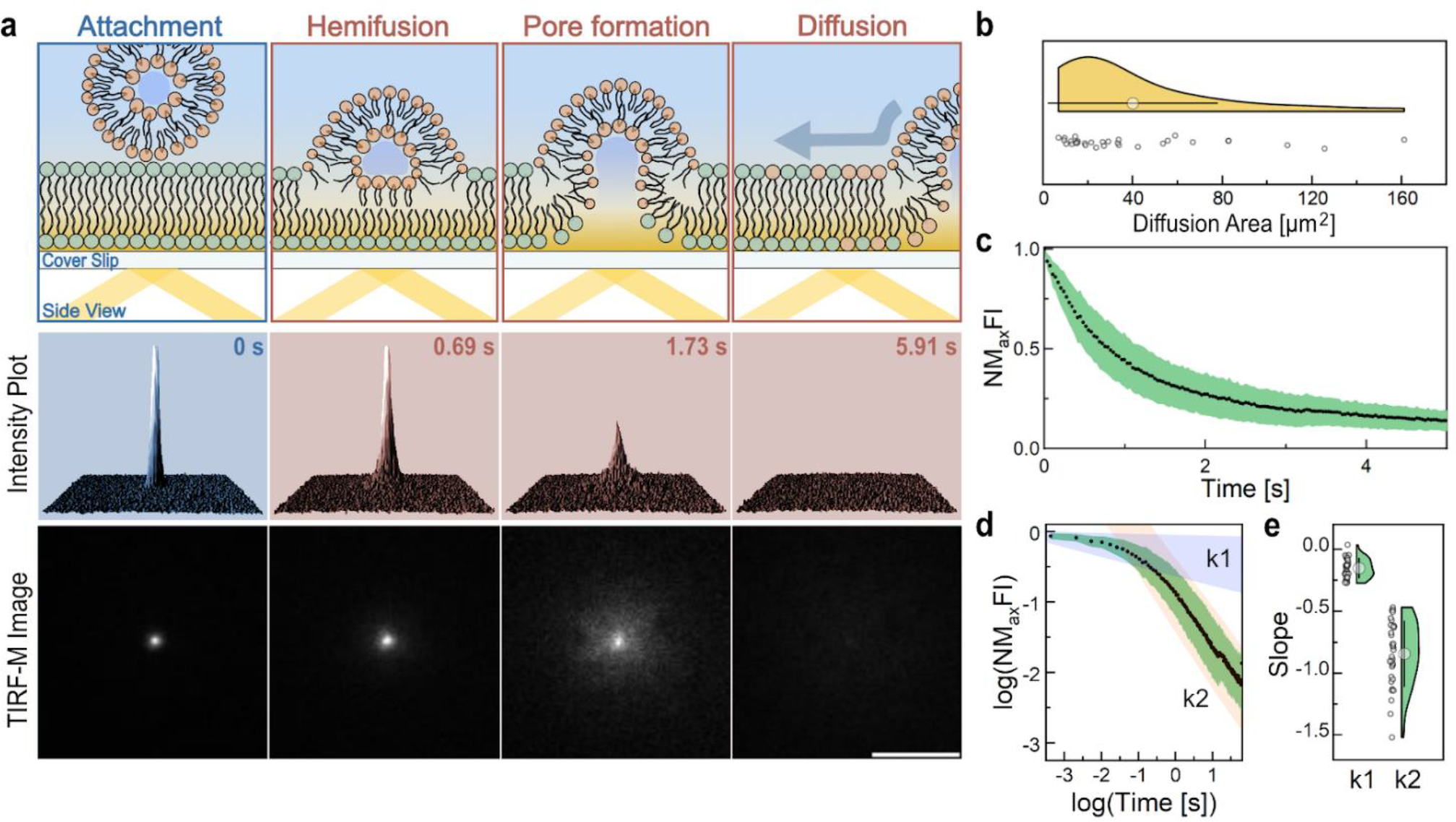
cFLs fuse with *E.coli* lipid bilayers. (a) Schematic depiction (top), three-dimensional intensity plot (middle) and corresponding TIRF images (bottom) of a representative time-lapse series displaying liposome attachment and lipid diffusion into the SLB. Scale bar, 10 µm. (b) Lipid diffusion area of single liposomes upon contact with SLB. Displaying Mean ± SD (Coef=1), *n* = 31. (c) Normalised maximum fluorescence intensity (NMFI) decay plot of cFLs upon attachment to SLB. Displaying Mean ± SD, *n* = 33. (d) Corresponding log-log plot suggesting a twostep membrane interaction with two distinct slope regimes in blue (k1) and orange (k2). Displaying Mean ± SD, *n* = 33. (e) Comparison of the interaction slopes k1 -0.15± 0.08 and k2 -0.84 ± 0.26 (Mean ± SD, *n* = 33).

To investigate the fusion kinetics, several single particle interactions were recorded and the change in maximum fluorescence intensity at the attachment site was calculated (Fig. 3c). The log-log plots of the decay curves revealed two distinct slope regimes of -0.15 ± 0.08 for k1 and -0.84 ± 0.26 for k2, corresponding to a two-step membrane interaction (Fig. 3d). Assumingly, the short living (∼100 ms) first slope is dominated by liposome attachment, while the second one shows the membrane merging and lipid diffusion process (Fig. 3d-e). Similarly, Dyett et al. reported a bi-phasic cubosome interaction with Gram-negative bacteria that follow comparable interaction slopes (k1 =∼ -0.16 and k2=∼ -1) but at prolonged time scales. According to their work, the first phase corresponds to liposome fusion with the OM followed by diffusion through the cell wall.^23^ Here, however, liposome fusion with the SLB system is completed within a few seconds, and the fusion process with a slope of -0.84 ±0.26 resembles a diffusion of a point source release in two dimensions as previously described by Dyett et al..^35^

### Lipids of cFLs are internalized by Gram-positive bacteria

By studying the interactions of liposomes with model systems like mica and the *E. coli* SLB, we demonstrated the ability of cFLs to form bilayers on anionic surfaces and to fuse with membranes, respectively. The interactions with living bacteria, however, may vary significantly. The envelope of Gram-positive bacteria is composed of an inner membrane that is encapsulated by a thick peptidoglycan network. Since this network does not have the same properties as a lipid bilayer, cFLs are not expected to fuse but could spread to form bilayers on the cell surface as demonstrated on a mica substrate.

Gram-positive *B. subtilis* cells were incubated with cFLs and imaged using SIM. Bacteria were identified by staining the cytoplasmic DNA (green, 488 nm) and the liposomes were visualized using Rh-PE labeling (orange, 561 nm). The reconstructed super-resolution microscopy images showed a mixture of Gram-positive bacteria that were targeted by the liposomes and others that remained untargeted (Fig. 4a). Neutrally charged control liposomes lacking DOTAP did not result in bacterial targeting (Supplementary Fig. 3). Therefore, electrostatic interaction is required to drive liposome attachment. As highlighted in Fig. 4a, the cFLs interacted heterogeneously with *B. subtilis* cells. The liposomes attached to the perimeter of the bacterium (highlighted as stars) or within the cytoplasm, overlapping with the cyto-plasmic DNA stain (highlighted with arrow, Fig. 4a-b). Additionally, the comparison between the fluorescence profile derived from the cytoplasmic detected Rh-PE signal and the fluorescence profile obtained through simulating a cytoplasmic staining revealed a similar profile (Fig. 4b). This evidence suggests liposome or single lipid internalization by *B. subtilis* cells, rather than cFLs enveloping the surface of the target cell. The uptake of small liposomes and fusion with the inner membrane might be achieved through the presence of large pores of up to 60 nm diameter that span the cell wall.^36^ Another possibility is the gradual uptake of attached lipids. Dyett et al. studied the interactions of cubosomes (cubic lipid nanoparticles) with Gram-positive *Staphylococcus aureus* cells proposing the possibility of lipid uptake by Gram-positive cells.^37^ To study how the fusogenic lipid DOPE affects the interaction with the cell envelope, control experiments with non-fusogenic liposomes (nFLs) composed of DOTAP and the structural lipid 1,2-dioleoylsn-glycero-3-phosphocholine (DOPC) were performed. As highlighted in Fig. 4c, nFLs lacking DOPE were able to target bacteria, however, they predominantly attached to the bacterial envelope (Fig. 4c). By comparing the Mander’s colocalization coefficient (MOC) (Supplementary information ‘Colocalization analysis’ section) between Rh-PE and the cytoplasmic DNA, we noticed a significantly stronger colocalization in presence of DOPE (Fig. 4d). Therefore, DOPE is essential for the interactions with the peptidoglycan network to drive lipid uptake by Gram-positive bacteria (Fig. 4e).

**Figure 4.**
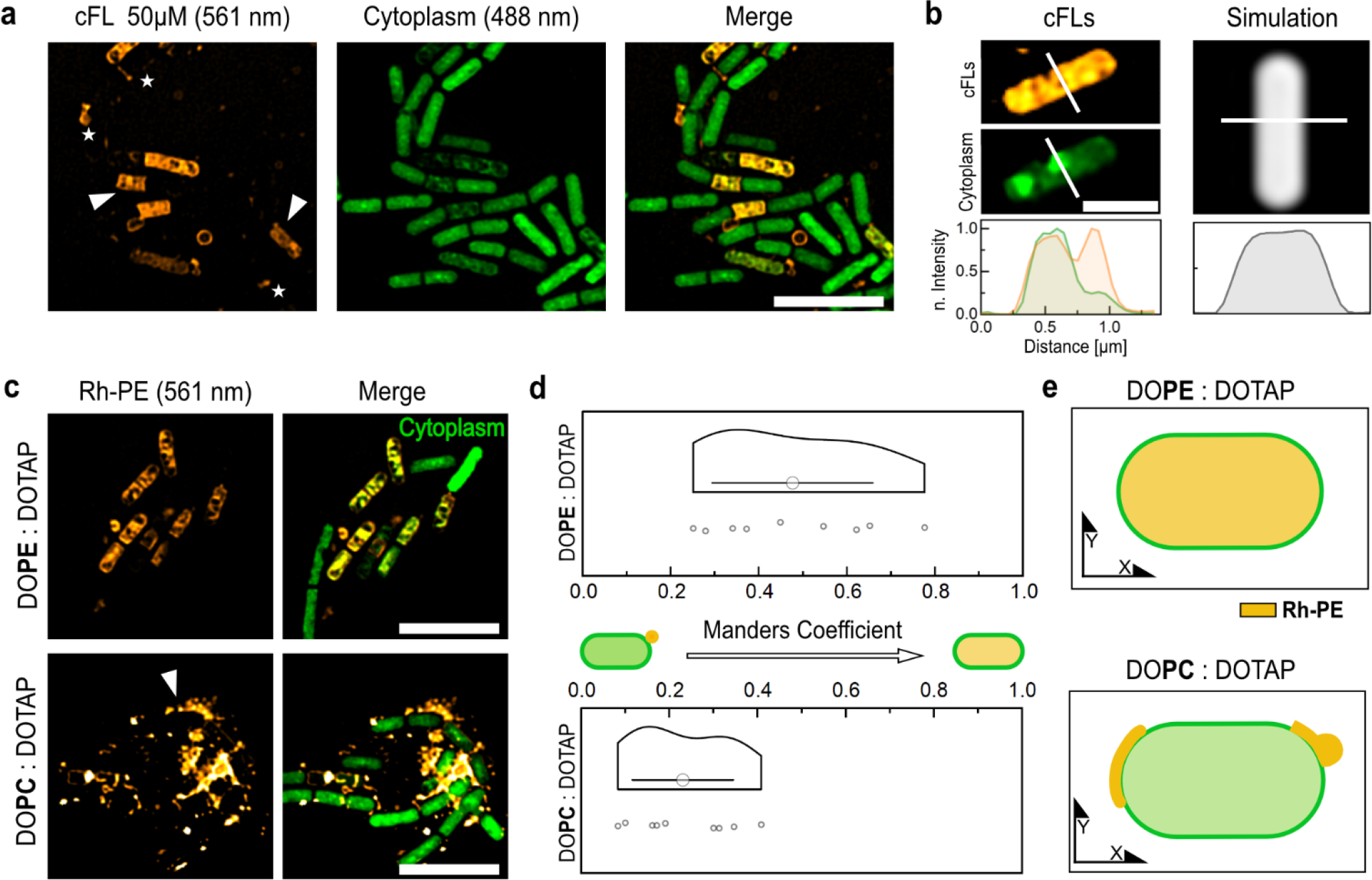
Super-resolution microscopy shows cFLs attachment and lipid internalization by Gram-positive *B. subtilis* cells. (a) SIM images of cFLs (orange) targeted *B. subtilis* cells (green). Rh-PE lipids were found to be attached to the cell perimeter (white star) or localized inside the cytoplasm (white arrow). Scale bar, 5 µm. (b) Representative SIM image and corresponding normalized fluorescence intensity plot (NFIP) of cFLs targeted bacteria in comparison to a cytoplasmic dye simulation. Scale bar, 1 µm. (c) Representative SIM images of *B. subtilis* cells incubated with cFL (DOPE:DOTAP, 1:1 mole ratio) and nFL (DOPC:DOTAP, 1:1 mole ratio) liposomes. Scale bar, 5 µm. White arrow highlights attached liposomes. (d) Mander’s colocalization coefficient (MOC) between Rh-PE and the cytoplasmic DNA demonstrates enhanced liposomal uptake in presence of DOPE. (DOPC:DOTAP n = 9, DOPC:DOPE: *n* = 9, means significantly different **). (e) Illustration of FL internalization and nFLs attachment of targeted Gram-positive *B. subtilis* cells.

### Cationic fusogenic liposomes merge with the outer membrane of Gram-negative bacteria

After confirming liposome fusion with a model membrane system and visualizing the lipid internalization by Gram-positive bacteria, we studied the interactions with Gram-negative bacteria. Specifically, cytoplasmic GFP-expressing BL21 cells (green, 488 nm) were exposed to cFLs (orange, 561 nm) and imaged using SIM. The reconstructed images showed a mixture of Gram-negative bacteria that were targeted by the cFLs and others that remained untargeted (Fig. 5a). The targeted cells exhibited a fluorescent halo in the 561 nm channel that surrounded the cytoplasmic signal (Fig. 5a,b). This distinct Rh-PE fluorescence profile of targeted cells agrees with the fluorescence profile obtained through simulating a bacterial membrane staining (Fig. 5b). The profiles demonstrate a fluorescence intensity maximum at the perimeter of the cell with the intensity gradually decreasing towards the midpoint (Fig. 5b). These results suggest the integration of cFLs into the OM of BL21 cells. Interestingly, in some bacteria, internal Rh-PE signals were observed along with a loss of GFP signal in the same area (Supplementary Fig. 4, Fig. 5a). Previous studies by Moreira et al. have reported increased cytoplasmic leakage when bacteria are targeted with fusogenic liposomes.^38^ Therefore, our observation could be a result of membrane destabilization and leakage of cytoplasmic GFP. However, as highlighted in Fig. 5a and Supplementary Fig. 4, the GFP signal is not uniformly decreased within targeted cells. Areas of decreased GFP signal within the cytoplasm may be caused by fluorescence resonance energy transfer (FRET) between GFP and internalized Rh-PE. This means that an excited GFP molecule transfers energy to Rh-PE, resulting in quenching of the GFP fluorescence signal in the same area as Rh-PE. To determine the effect of the fusogenic lipid DOPE, control experiments with the previously introduced nFLs were performed. Cells targeted by nFLs showed segmented halo structures (Fig, 5c), which either suggest nFL attachment and spreading along the cell perimeter or partial membrane integration. Due to the segmented Rh-PE staining, we analyzed the degree of coverage (see ‘Colocalization analysis’ section) upon liposome exposure. The cFLs samples revealed two populations, one displaying a low degree of coverage which indicates liposome attachment and a higher degree of coverage which suggests liposome uptake into the OM (Fig. 5d). nFLs, however, showed a significantly bigger population of low degree of coverage (Fig. 5d). DOPC, as a lamellar phase promoting lipid with a spontaneous curvature close to zero,^28^ stabilizes lipid bilayers but is known to greatly reduce the transfection efficiency of lipoplexes.^6^ Nevertheless, Laune et al. have demonstrated the fusion ability of DOPC containing liposomes with high (75 mol%) DOTAP content.^25^ Hence, the segmented Rh-PE staining at the cell perimeter may result from reduced membrane uptake ability of the nFLs or liposomal attachment to the envelope (Fig. 5e). These results demonstrate the importance of DOPE to drive efficient liposome fusion with the OM of *E. coli* BL21 cells (Fig. 5e).

**Figure 5.**
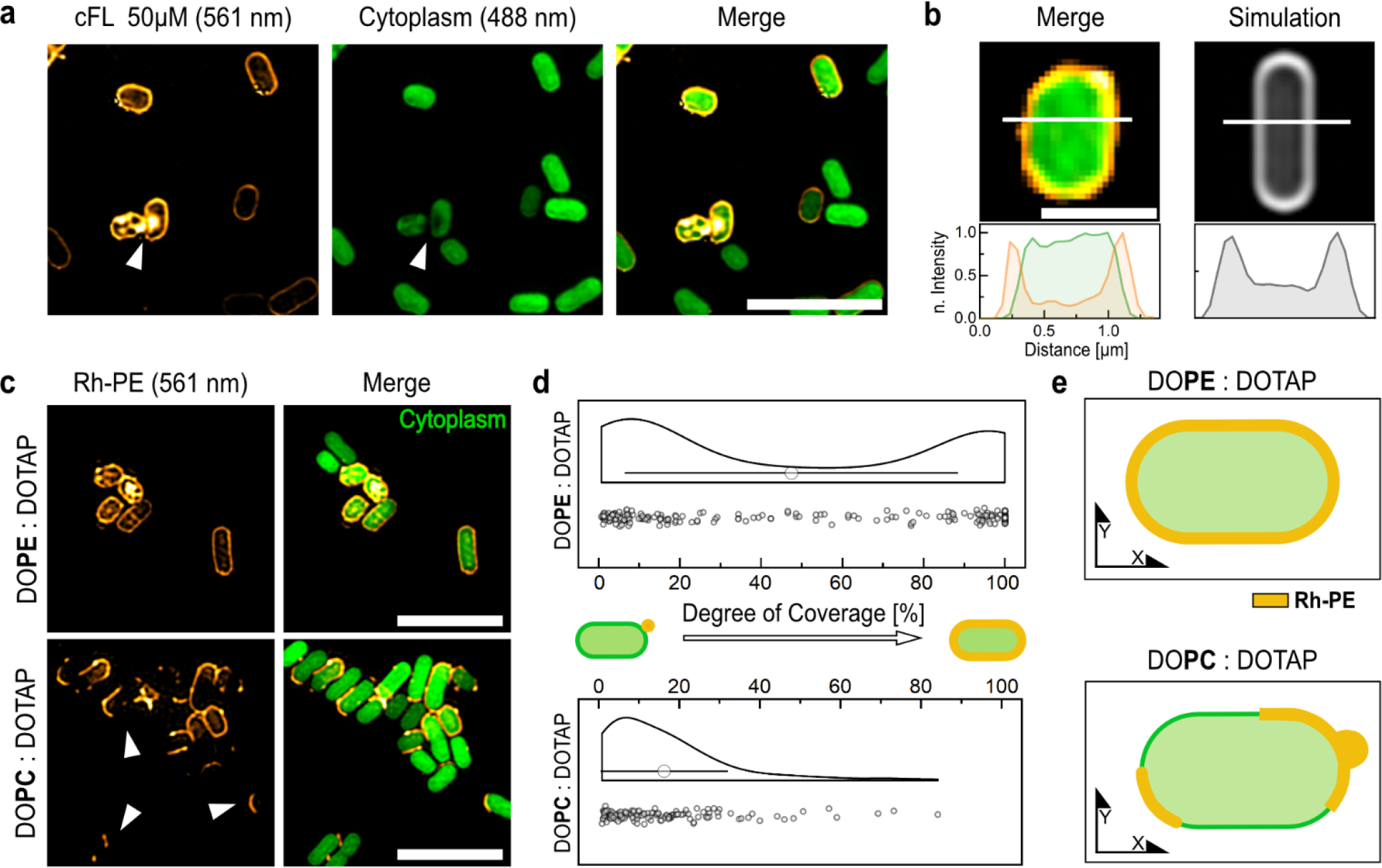
Super-resolution microscopy shows the integration of cFLs into the OM of Gram-negative BL21 cells. (a) SIM images of cFLs (orange) targeted *E.coli* BL21 cells. Scale bar = 5 µm. cFLs targeted cells displayed a halo signal around the cytoplasmic GFP signal (green). White arrow indicates Rh-PE signal within cytoplasm with lack in GFP signal at the same site. (b) Representative SIM image and corresponding NFIP of cFLs targeted bacteria in comparison to a membrane dye simulation. Scale bar = 1 µm. (c) Representative SIM images of BL21 cells against cFL (DOTAP:DOPE, 1:1 mole ratio) and nFL (DOPC:DOTAP, 1:1 mole ratio) liposomes. Scale bar = 5 µm. (d) Degree of coverage of cells targeted by cFLs and nFLs demon strates a population of full membrane integration (100 % coverage) in presence of DOPE and a population of low degree coverage, probably referring to attachment and partial lipid integration, which is in turn equally observed in the nFL control. (DOPC:DOTAP n = 127 individual bacteria, DOPC:DOPE: *n* = 215 individual bacteria, means are significantly different ****). (e) Illustration of full cFL uptake into the OM of Gram-negative cells against the partial integration and attachment of nFLs.

### Heterogeneous interaction dynamics with Gram-negative bacterial envelopes

Having demonstrated the integration of cFLs into the OM of *E. coli* cells using SIM, we sought to understand the real-time interaction dynamics at both the single particle and single cell level. Employing highly inclined and laminated optical sheet microscopy (HILO-M)^39^ the interactions of cFLs with Poly-L-Lysine (PLL) immobilised BL21 cells were recorded (Fig. 6a-b). The time-lapse images revealed three distinct interactions upon membrane encounter (Fig. 6c): the attachment of liposomes to the perimeter of a bacterium without recording a change in the fluorescence intensity (Fig. 6ci), attachment of liposomes followed by Rh-PE diffusion into the OM with a fluorescence intensity decay of less than one half of the peak attachment intensity (>50 % of initial fluorescence) (Fig. 6cii), and complete liposome integration with Rh-PE diffusion throughout the entire OM without leaving high residual fluorescence signal at the attachment site (<50 % of initial fluorescence) (Fig. 6c iii). We assign these heterogeneous interactions to liposome attachment, hemifusion, and fusion (Fig. 6c). Videos of the fusion events depicted in Fig. 6c are available in Supplementary Videos 2-4. The absence of membrane fusion following liposomal attachment could be attributed to insufficient membrane interactions. The thick lipopolysaccharide (LPS) enveloping the outer membrane of BL21 cells may present a barrier to membrane engagement, thereby impeding lipid mixing and membrane fusion. If transitioning to the hemifusion state, the inner leaflets of cFL and the bacterial OM remain separated, leaving behind a high residual Rh-PE fluorescence at the attachment site. This hemifusion intermediate is attributed to the presence of the hexagonal phase-promoting lipid DOPE.^28^ While cFL attachment events exhibited constant fluorescence signals over extended periods (Fig. 6d, green), hemifusion and fusion events (Fig. 6e, orange) resembled the interactions found for cFLs against SLBs, with the fluorescence signal at the attachment site gradually decayed due to lipid diffusion into the encountered membrane. The normalised maximum fluorescence intensity (NMFI) decay curves of hemifusion and fusion events were fitted with an exponential function (I = exp(-λt) + A; where I describes the maximum fluorescence intensity and t is time) to extract the lipid diffusion exponent (λ) and the fluorescence plateau (A). Surprisingly, the mean lipid diffusion exponents measured for BL21 (17.7 ± 4.24) cells were larger than the fusion exponents measured for SLBs (1.31 ± 0.14) (Fig. 6e). Even though the OM of Gram-negative bacteria is a complex membrane with high protein content, liposome fusion with a SLB might be slower because of strong interaction with the glass support.^40^ For instance, Przybylo et al. reported significantly reduced lipid diffusion in an SLB compared to a giant vesicles model.^41^ In order to distinguish between hemifusion (>50 % of initial fluorescence) and fusion (<50 % of initial fluorescence) events, the residual fluorescence intensity values (plateau, A) at the attachment site were compared (Fig. 6f). We found full fusion of cFLs with SLBs, while 78.6% of liposome interactions with E. coli BL21 cells resulted in hemifusion events. This suggests that the OM with its LPS cover effectively acts as a barrier, preventing liposome fusion and direct cargo release into the periplasmic space.

**Figure 6.**
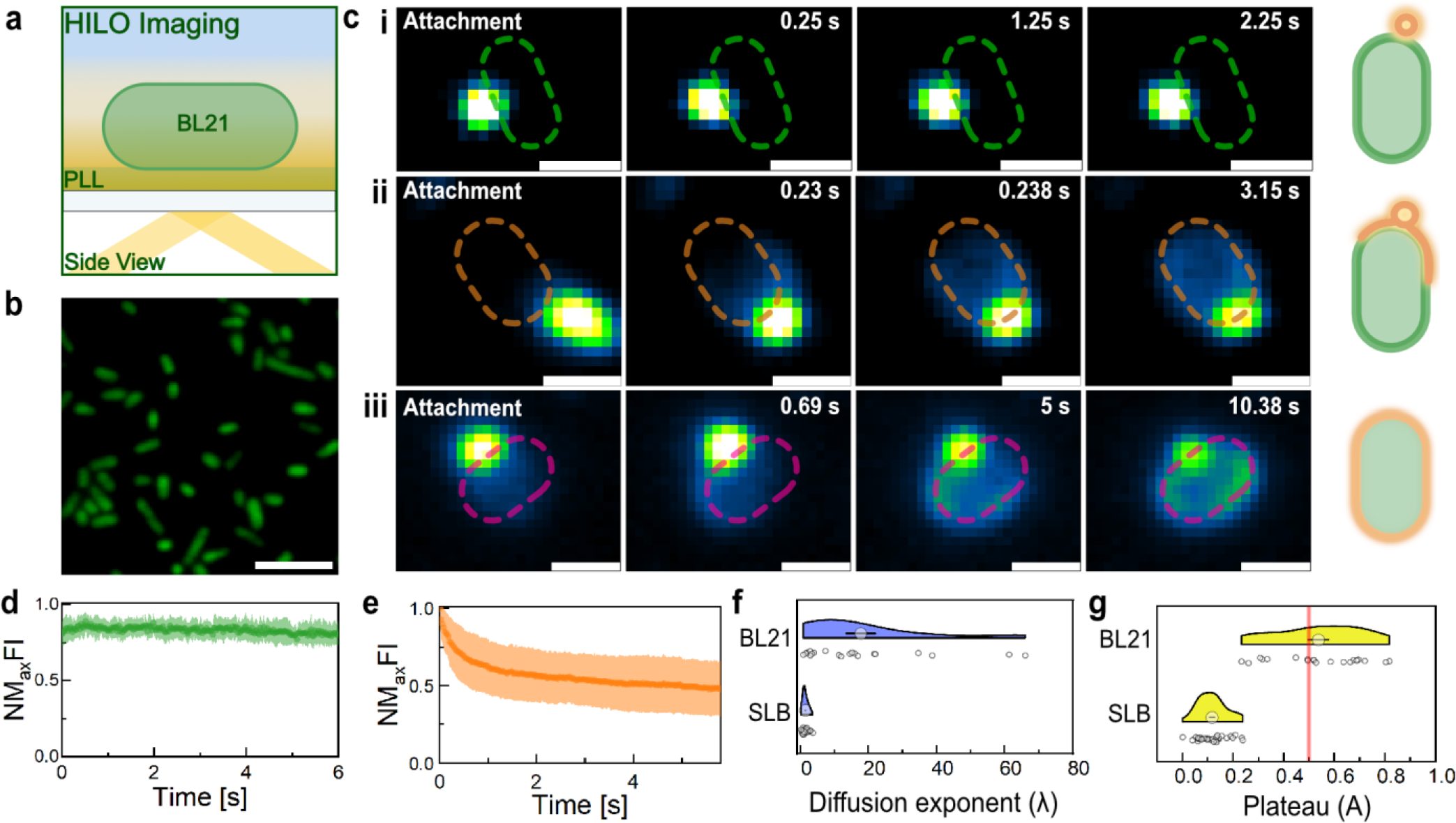
cFLs show heterogeneous interaction dynamics with OM of Gram-negative bacteria. (a) Schematic illustration of HILO-M on cytoplasmatic GFP expressing BL21 cells immobilized on glass coverslips using PLL. (b) Representative HILO image of PLL immobilized BL21 bacteria before the addition of cFLs. (c) Representative HILO image time-lapses of three interaction phenomena recorded for cFLs against BL21 cells. Scale bar, 1 µm. (right) Illustration of the assigned cFLs interaction types: attachment, hemifusion, and fusion. (d-e) NMFI plots of targeted bacteria, displaying liposome attachment (d, green, n=12) and fusion (e, orange, n=15) events. Results displayed as Mean ± SD. (f-g) Lipid diffusion exponent (λ) and normalized fluorescence plateau (A) plots extracted from NMFI decay curves of cFLs interactions with SLB(Mean ± SEM, n=32) and BL21 cells (Mean ± SEM, n=20). Red line highlights plateau cutoff to distinguish hemi (<50 % of initial fluorescence) and full fusion (>50 % of initial fluorescence) events.

### Compromised membrane integrity upon cFL encounter facilitates antibiotic targeting

The densely packed LPS layer acts as a permeability barrier that allows small molecules of less than ∼600 Da to enter the periplasmic space.^42^ However, antibiotics with higher molecular weight and low polarity are hindered from reaching intracellular targets. As a result, the usage of several Gram-positive-only drugs as broad-spectrum antibiotics is limited.^43^ The integration of artificial lipids with positive charge and high curvature may, however, compromise the outer membrane barrier function, facilitating the targeting ability of Gram-positive-only antibiotics.

While multiple liposomes attached and fused with individual Gram-negative bacteria (Supplementary Figure 5), an excess of charged and curved lipids is integrated into the outer membrane. To assess the effect of this artificial lipid integration on bacterial activity, an 3-(4,5-dimethylthiazol-2-yl)-5-(3-carboxymethoxyphenyl)-2-(4-sulfophenyl)-2H-tetrazolium (MTS) assay was performed. The cell vitality of BL21 cells was significantly reduced at cFLs concentrations above 10 µM (Fig. 7a). In addition to these population assays, super-resolution microscopy imaging was used to visualize the temporal effects of lipid integration on the morphology of the Gram-negative outer membrane. Some SIM images of liposome-fused bacteria showed detached Rh-PE halo signals, suggesting that bacteria were trapped within a large liposomal bubble (Fig. 7b). This observation may be the result of excessive lipid packaging into the OM, which generates lateral stress and promotes membrane detachment from the underlying peptidoglycan cell wall (Fig. 7c). Moreover, time-lapse images of targeted bacteria in PBS showed membrane vesicle formations at the cell surface (Fig. 7d). Interestingly, the accumulation of phospholipids, such as phosphatidylethanolamine, from the inner membrane into the outer leaflet of the OM has been proposed as one mechanism for outer membrane vesicle formation and increased membrane permeability.^11,44,45^ Therefore, integration of DOTAP and DOPE could potentially cause a breach in OM integrity, resulting in packing disruptions at the interface of LSP and phospholipids and cause membrane detachment. These results are further supported by Moreira et al. observation of increased cytoplasmic leakage upon fusogenic liposome encounter.^38^ Therefore, we sought to test if the co-delivery of macromolecular antibiotics, such as the fluorescently labeled antibiotic Vancomycin-BODIPY, is enhanced through the permeabilization of the bacterial membrane. BL21 cells co-incubated with cFLs and Vancomycin-BODIPY (∼1.4 kDa) were imaged using SIM. The reconstructed images and the mean Spearman coefficient of ∼0.5 describe a high degree of colocalization between liposome-targeted and Vancomycin targeted cells (Fig. 7e-f). In contrast, control experiments without cFLs or Vancomycin exposure exhibited low spearman coefficients of ∼0.1 (Fig. 7f). Therefore, FLs disturb the bacterial envelope, making bacteria more susceptible to large antibiotics that otherwise would not be able to penetrate through the OM.

**Figure 7.**
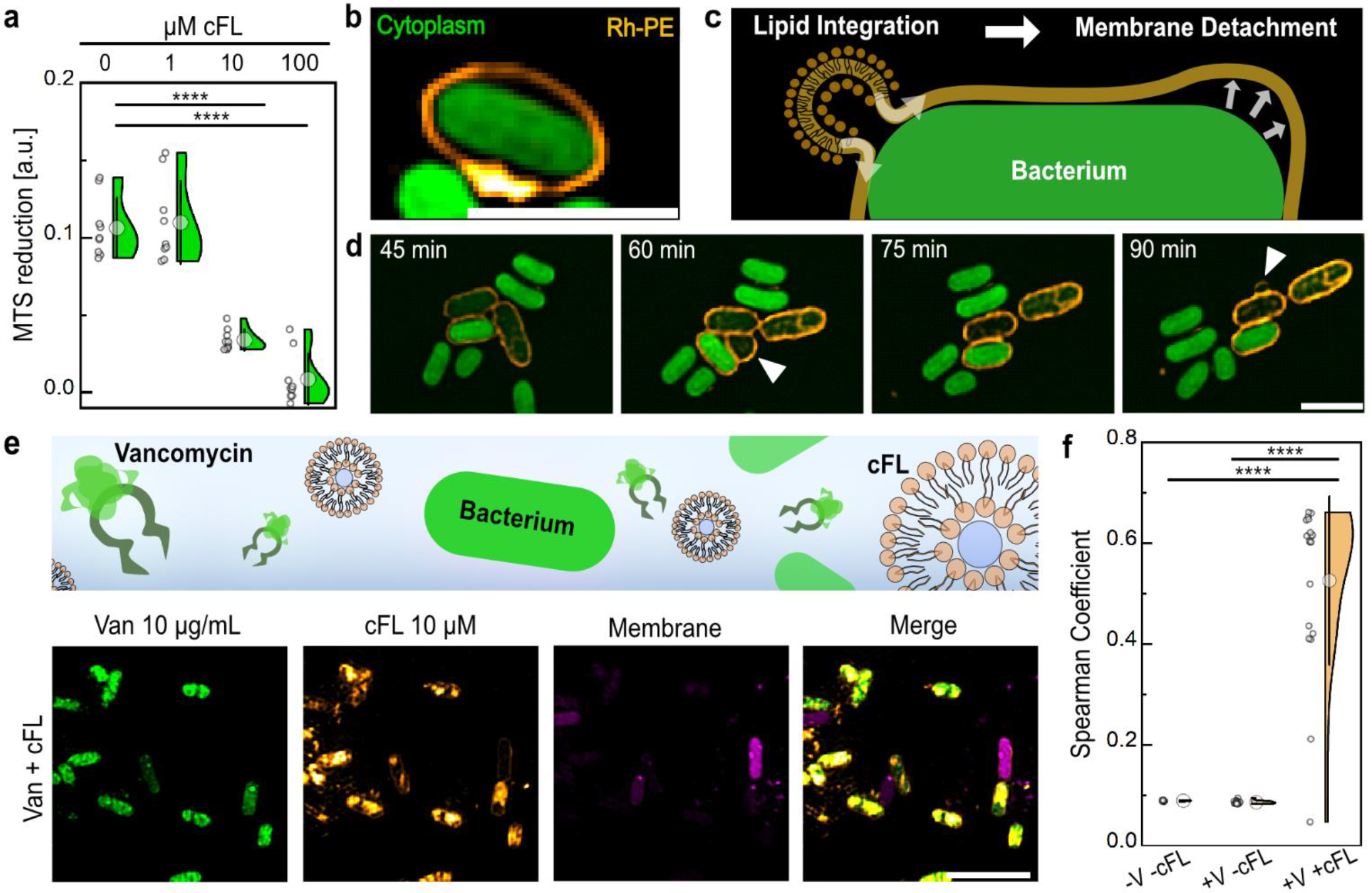
cFLs impair OM integrity and enhance antibiotic co-delivery. (a) Colorimetric cell viability assay measuring 3-(4,5-dimethylthiazol-2-yl)-5-(3-carboxymethoxyphenyl)-2-(4-sulfophenyl)-2H-tetrazolium (MTS) reduction in presence and absence of fusogenic liposomes. BL21 cell vitality is significantly reduced in presence of 10 µM and 100 µM cFLs. (b) Representative SIM image of a BL21 cell (green) surrounded by a detached Rh-PE halo signal (orange). Scale bar, 2 µm. (c) Schematic illustration of OM detachment driven by excessive cFLs lipid integration. (d) SIM time-lapse series of cFLs (orange) targeted BL21 cells in PBS. White arrows indicate the appearance of membrane vesicles. (e-f) Vancomycin (green) targeting of BL21 cells upon addition of cFLs (orange). Cells were exposed to a mixture of vancomycin and cFLs before being imaged using SIM. (e) Representative SIM images of vancomycin and cFLs target BL21 cells. Scale bar, 5 μm. (f) Colocalization between Vancomycin and cFLs using the Spearman coefficient method. (Mean ± SD).

Besides affecting membrane integrity, we observed bacterial agglomeration in presence of cFLs (Supplementary Fig. 6). These bacterial clumps, forming through electrostatic interactions, were visible to the naked eye (Supplementary Fig. 6) and could lead to severe complications if induced in the circulatory system. Furthermore, introducing charged particles into biological systems may drive the attachment of negatively charged molecules that form a protective corona that changes their biophysical properties and therefore limits their targeting abilities.^35^

## CONCLUSIONS

We used super resolution microscopy to gain valuable insights into the heterogeneous interactions between fusogenic liposomes and bacterial envelopes, with important implications for the development of the next generation of antibiotic delivery systems. The liposomes used in this study facilitated bacterial targeting through cationic lipids, while structural components were used to mediate membrane fusogenicity. By using SIM, lipid internalisation by Gram-positive bacteria was demonstrated. The interactions with Gram-negative bacteria were heterogeneous, showing liposome attachment, hemifusion and full liposome integration into the outer membrane. The number of recorded hemifusion events increased with increasing membrane complexity. Hence, generalising liposome interactions from one bacterium to another may be inaccurate, emphasising the need to study liposome attachment and fusion at the single cell level and with pathogenic bacteria, which frequently exhibit more complex envelope structures. To enhance fusion pore formation and facilitate content delivery across the outer membrane of Gram-negative bacteria, the inclusion of shape-changing lipids, also known as malleable lipids,^29^ might be a viable strategy. However, we have shown that the incorporation of artificial lipids into the outer membrane caused lipid packaging disruption, vesiculation, and a loss of membrane barrier function. By codelivering antibiotics and fusogenic liposomes, large antibiotics that cannot penetrate the outer membrane can now reach their intracellular target. This ability of fusogenic liposomes to modify bacterial envelope characteristics presents a promising tool to fight antimicrobial resistance even in the absence of antibiotic agents. The targeting ability of the cationic liposomes used in this study, however, is limited by the formation of a protein corona and non-specific bacterial targeting. To address these issues, the incorporation of targeting moieties such as lectins,^46^ aptamers,^47^ or DNA zippers ^48,49^ through chemical conjugation can enable specific targeting of pathogenic bacteria, enhancing their therapeutic index. Additionally, fusogenic liposomes may show exciting prospects in cancer research to insert immune cell activating components and altering membrane characteristics to enhance drug susceptibility. Overall, we aim to emphasize the significance of comprehending the interactions between nanocarriers and their targets. Improving our understanding of these interactions is not only essential to finding new ways to fight infections but also for advancing various aspects of biomedical research.

## ASSOCIATED CONTENT

The Supporting Information is available free of charge at [link]. Methodology and additional experimental details include: AFM images of E. coli total lipid extract SLB, TIRF-M time-lapse images of cFL interactions with glass supports, SIM images of BL21 and *B. subtilis* cells exposed to cFLs and nFLs, SIM image of BL21 cells highlighting the internal Rh-PE signal accompanied by a loss of GFP fluorescence, multiple interactions of cFLs with individual bacteria, charge-driven bacterial agglomeration, SIM of Vancomycin-targeted BL21 cells, image processing pipelines, and descriptions for Video S1–S4.

Video S1 (AVI)

Video S2 (AVI)

Video S3 (AVI)

Video S4 (AVI)

## AUTHOR INFORMATION

### Author Contributions

The manuscript was written through contributions of all authors and all authors have given approval to the final version of the manuscript.

## ACKNOWLEDGMENT

The authors would like to thank Professor Ulrich F. Keyser and Dr. Lorenzo Di Michele for their helpful discussions on the project.

## ABBREVIATIONS

A: fluorescence plateau
AFM: Atomic force microscopy
cFLs: Cationic fusogenic liposomes
DLS: Dynamic light scattering
DOPE: 1,2-dioleoyl-sn-glycero-3-phosphoethanolamine DOPC, 1,2-dioleoyl-sn-glycero-3-phosphocholine
DOTAP: 1,2-dioleoyl-3-trimethylammonium-propane
FLs: Fusogenic liposomes
FRET: Fluorescence resonance energy transfer
HILO-M: Highly inclined and laminated optical sheet microscopy
λ: diffusion exponent
LPS: Lipopolysaccharide
MOC: Mander’s colocalization coefficient
MTS: 3-(4,5-dimethylthiazol-2-yl)-5-(3-carboxymethoxyphenyl)-2-(4-sulfophenyl)-2H-tetrazolium
nFLs: Non-fusogenic liposomes
NMFI: Normalized maximum fluorescence intensity
OM: Outer membrane
PLL: Poly-L-Lysine
Rh-PE: 1,2-dioleoyl-sn-glycero-3-phosphoethanolamine-N-(lissamine rhodamine B sulfonyl)
SIM: Structured illumination microscopy
SLB: Supported lipid bilayer
TEM: Transmission electron microscopy
TIRF-M: Total internal reflection microscopy

---

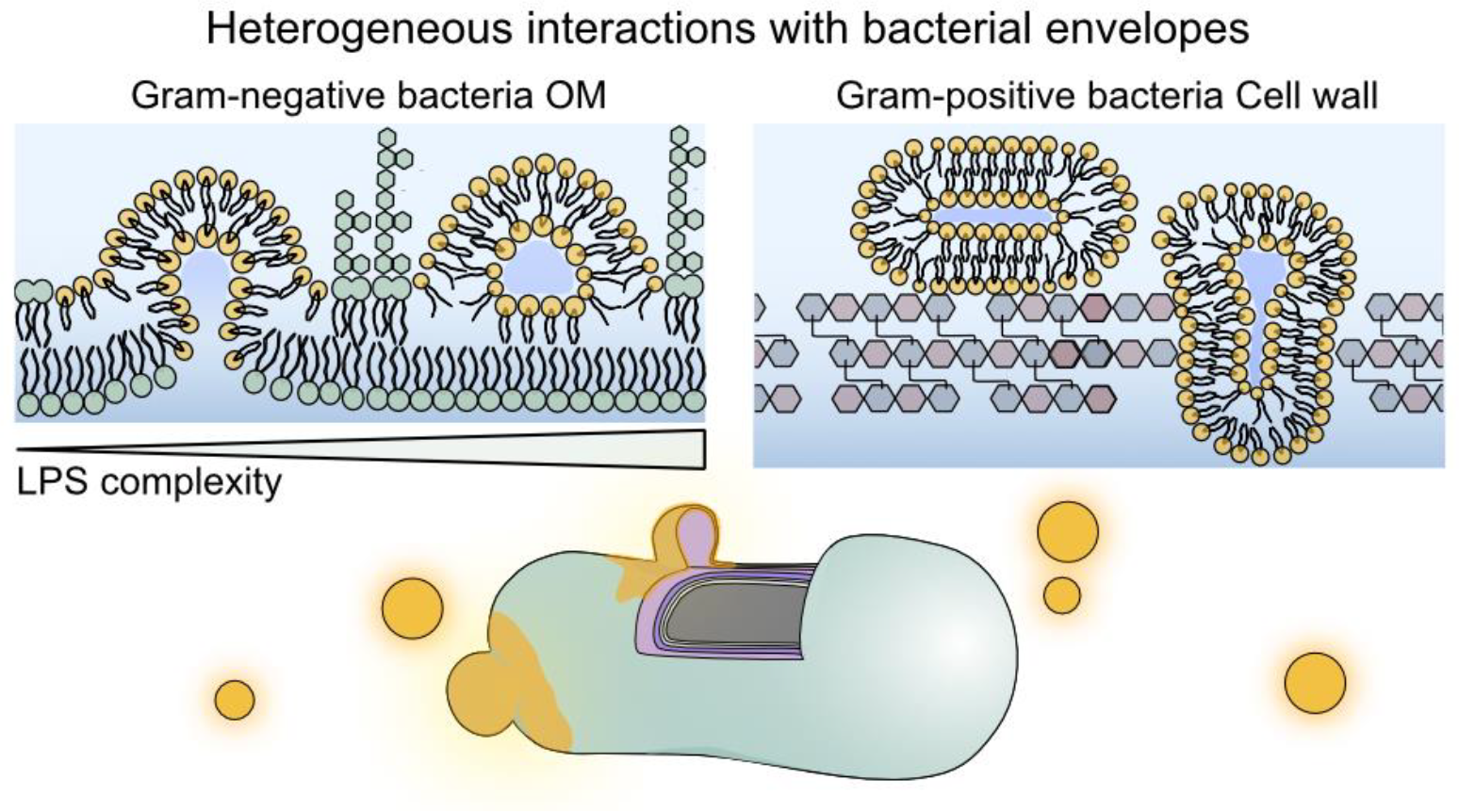

