## Supporting Information for "Molecular mechanisms of liposome interactions with bacterial envelopes"

---

**Table of Contents**

**Experimental section** **3**

Liposome preparation 3

Dynamic light scattering (DLS) and zeta potential measurements 3

Bacterial culture and staining 4

Supported lipid bilayer formation 4

Microscopy 4

Atomic force microscopy 4

Structured Illumination Microscopy 5

Total internal reflection fluorescence Microscopy 5

3-(4,5-dimethylthiazol-2-yl)-5-(3-carboxymethoxyphenyl)-2-(4-sulfophenyl)-2H-tetrazolium (MTS) assay 6

Data analysis 6

TIRF-M analysis of Fusion Events 6

Colocalization analysis 7

Mander's colocalization analysis between cytoplasmic DNA and Rh-PE 7

Spearman colocalization analysis between Vancomycin-BODIPY and Rh-PE 7
Degree of coverage analysis between Gram-negative Bacteria and cFLs 7
Modelled SIM data 7
Data visualization and statistical analysis 8
**Supplementary figures** **9**
**Bibliography** **18**

### Experimental section

#### Liposome preparation

Small vesicles were prepared with the thin film hydration technique, as previously reported.[1] 1,2-dioleoyl-3-trimethylammonium-propane (chloride salt) (DOTAP), 1,2-dioleoyl-sn-glycero-3-phosphoethanolamine (DOPE), 1,2-dioleoyl-sn-glycero-3-phosphocholine (DOPC), and 1,2-dioleoyl-sn-glycero-3-phosphoethanolamine-N-(lissamine rhodamine B sulfonyl) (ammonium salt) (Rh-PE) reconstituted in chloroform were purchased from Avanti Polar Lipids and stored at -20ºC. Cationic fusogenic liposomes (cFL) were mixed in a glass vial from a 1:1 mole ratio of DOPE:DOTAP while non-fusogenic cationic liposomes (nFL) were prepared from a 1:1 mole ratio of DOPC:DOTAP. For fluorescent labeling at 561 nm excitation, 0.25 mole % Rh-PE was added to the corresponding lipid mixture. The chloroform was evaporated using a nitrogen stream and the dried lipid film was further desiccated for 2-3 hours under vacuum. The thin lipid film was hydrated in 1 mL of 1X PBS for 30 minutes at room temperature resulting in a final lipid concentration of 10 mM. Single unilamellar vesicles were prepared through extrusion following the producer's protocol with a Avanti Mini Extruder (Avanti® Polar Lipids). Whatman® Filter Supports and Whatman® Nuclepore Track-Etched Membranes (100 nm) were obtained from Sigma Aldrich. The samples were stored for up to two weeks at 4°C.

#### Dynamic light scattering (DLS) and zeta potential measurements

The average hydrodynamic diameter and zeta (ξ) potential of the liposomes were measured using a Zetasizer Nano ZSP (Malvern Panalytical) with an excitation wavelength of 633 nm and with a fixed scattering angle of 173°. "Hydrodynamic size distribution measurements were conducted on 1 mL of liposomes with a concentration of 500 µM. Each sample was measured three times, with each measurement consisting of at least 12 runs. The output data included the average particle size distribution (Z average (d.nm)) and the polydispersity index (PDI), which measures the heterogeneity of the particle size distribution. Data are presented as mean ± standard deviation and were calculated from a number of repeats of three independent experiments.

Zeta potential measurements were performed on 1 mL of samples using DTS1070 cells. Liposomes were diluted in 1X PBS to a final concentration of 500 uM. The overnight culture of bacteria was 1:100 diluted in fresh LB media and grown to an OD600 of 0.5 before being washed three times through centrifugation at 6.000 rpm and resuspended in 1X PBS. Each sample was measured three times, with each measurement consisting of at least 12 runs. Data are presented as mean ± standard deviation and were calculated from a number of repeats of three independent experiments.

**Bacterial culture and staining**

10 mL LB medium was inoculated with either *E. coli* BL21 (DE3) (Invitrogen) or *B. subtilis* (BS168) (Invitrogen) cells and grown overnight in a shaking incubator at 37ºC. The starter cultures were 1:100 diluted in fresh LB medium and further grown until reaching an OD600 of 0.5. The culture was then washed three times through centrifugation (6000 rpm, 2min) and resuspension in 1X PBS. The *E. coli* cells used for cFLs interactions were either fluorescently labeled at 488 nm excitation by expression of a cytoplasmic GFP protein (pUC19GFP plasmid) or membrane stained with the membrane stain MitoTracker™ Deep Red FM (Thermo Fisher). *B. subtilis* cells were fluorescently live labelled with the cytoplasmic SYTO™ 9 Green Fluorescent Nucleic Acid Stain (Thermo Fisher). For the dye labelling procedure the freshly washed cells were incubated with either 5 µM SYTO™ 9 or 1 µM MitoTracker™ Deep Red for 15 minutes in a shaking incubator at 37ºC. The cells were washed three times as previously described before being used for further experiments.

**Supported lipid bilayer formation**

Small vesicles for the formation of Supported Lipid Bilayers (SLBs) were prepared using the thin film hydration technique, as previously described.[1] *E. coli* total lipid extract and 1,2-dioleoyl-sn-glycero-3-phosphoethanolamine-N-(7-nitro-2-1,3-benzoxadiazol-4-yl) (ammonium salt) (NBD-PE), reconstituted in chloroform, were purchased from Avanti Polar Lipids and stored at -20ºC. For fluorescent labeling at 488 nm excitation, 0.25 mole % NBD-PE was added to the *E. coli* Total Lipid Extract. The liposomes were prepared as previously described (see "Liposome preparation" section), resulting in liposomes in 1X PBS with a lipid concentration of 4 mg/mL. The samples were stored at 4°C for up to two weeks.

For the formation of the SLBs, glass coverslips (Academy, 22x40 mm, 0.16-0.19 mm thick) that were cleaned with acetone and isopropanol or µ-slide 8 well high glass bottom (Ibidi) were incubated with 200 µL of 2 mg/mL E. coli liposomes for 30 minutes under continuous shaking at room temperature. The samples were then washed three times with 1x PBS. The procedure was repeated once to fill holes in the SLB. The bilayer was allowed to rest for approximately 30 minutes after washing to facilitate healing from potential disruptions during the washing procedure. AFM images were acquired to confirm the formation of the SLB (see Supplementary Figure 1).

**Microscopy**

### Atomic force microscopy

Atomic force microscopy (AFM) images of cFLs and *E. coli* SLBs were acquired using a BioScope Resolve microscope (Bruker) in scanasyst mode, employing Peakforce HIRS-FA probes (Bruker) with a nominal spring constant of 0.35 N m−1 and a resonant frequency of 165 kHz. cFLs at a concentration of 10 nM were deposited onto freshly cleaved mica substrates and incubated for 10 minutes before being washed with 3 mL of 1X PBS. AFM images were recorded at scan speeds of 1 Hz, with tip-sample interaction forces ranging between 100 and 300 pN.The recorded topography data were first-order flattened using Nanoscope analysis software 2.0 (Bruker) before height measurements on the bilayers were conducted by capturing cross-sections across various regions of interest. To quantify the liposome spreading area, AFM images were converted to 8-bit images using Fiji (ImageJ) and then thresholded to select the bilayer patches. The liposome spreading area was determined using the built-in particle analyzer. The image processing pipeline is illustrated in Supplementary Fig. 7.

### Structured Illumination Microscopy

1 mL of a bacterial suspension (OD600 of 0.5) was incubated with liposomes at varying concentrations, as indicated in the main text, in a shaking incubator at 37ºC for 15 minutes. For co-delivery experiments involving Vancomycin and BODIPY™ FL Conjugate (Invitrogen), bacteria were incubated with 10 µM cFLs and 10 µg/mL Vancomycin in 1X PBS. After the incubation, the cell suspensions were vortexed, centrifuged (6000 rpm, 2 minutes), and then resuspended in 1x PBS. The bacteria were deposited (1 µL) onto a glass slide (Academy, 22x40 mm, 0.16-0.19 mm thick) and immobilized beneath a custom-made 5 mm x 5 mm agarose pad (1% w/v). The agarose pad was covered with a glass coverslip to prevent drying before imaging.

Fluorescence microscopy images were acquired using a 3-colour structured illumination microscopy (SIM) technique.[2] A 60x 1.2 NA water immersion lens (UPLSAPO 60XW, Olympus) focused the structured illumination pattern onto the sample and captured the emitted fluorescence light, which was then projected onto an sCMOS camera (Orca-flash 4.0, Hamamatsu). The excitation wavelengths used were 488 nm (iBEAM-SMART-488, Toptica) for bacteria, 561 nm (OBIS 561, Coherent) for Rh-PE labeled liposomes, and 640 nm (MLD 640, Cobolt) for membrane-stained bacteria. Image acquisition was performed using previously described custom SIM software.[3] SIM reconstructions were carried out using the open-source software FairSIM,[4] following best practices for parameter selection.[5] Each pixel measured 107 nm during acquisition, corresponding to 53.5 nm after reconstruction.

### Total internal reflection fluorescence Microscopy

Time-lapse Total Internal Reflection Fluorescence (TIRF) microscopy images were acquired for *E. coli* Supported Lipid Bilayers (SLBs) and whole bacteria exposed to cFLs. The SLBs were prepared in Ibidi wells, as previously reported in "Supported Lipid Bilayer formation". The E. coli cells were immobilized on 0.1% (w/v) poly-L-lysine (PLL) coated Ibidi wells. The Ibidi wells were subjected to O2 plasma treatment before the PLL suspension was incubated for 15 minutes. The surface was then washed with deionized water (DI H2O) and dried using continuous nitrogen flow. GFP-labeled bacteria were prepared in 1X PBS at an optical density (OD) of 0.5 and incubated on the PLL-coated glass surface for 10 minutes before being washed with 1X PBS. Subsequently, 200 µL of 1X PBS was added to either the *E. coli* SLB-coated wells or BL21 cell-immobilized wells. Initially, the 488 nm excitation was used to focus the SLB or bacteria into the imaging plane, following which the 561 nm excitation channel was used to record the addition of 100 nM cFLs.

Time-lapse TIRF-M experiments involving liposome fusion were conducted using a custom-built microscope based on a microscope frame (IX-73, Olympus), equipped with a 100x 1.49 NA oil objective lens (UAPON100XOTIRF, Olympus). The excitation wavelengths employed were 488 nm (Sapphire LDP, Coherent) for SLBs or bacteria and 561 nm (Jive 05-01, Cobolt) for Rh-PE labeled liposomes. The samples were imaged in TIRF mode, and the fluorescence was captured using an electron-multiplying CCD (iXon Ultra, Andor). The pixel size was measured to be 117 nm. TIRF recordings were acquired at an exposure time of 34 ms for SLB experiments using an EM gain of 200 over a 512x512 pixel region.

TIRF experiments involving bacteria were conducted with exposure times of 10 ms over a 256x256 pixel region or 22 ms over a 512x512 pixel region.

#### Cell vitality assay

E. coli cells resuspended in PBS were prepared as previously described. 200 µL of the bacteria was transferred into flat-bottomed 96-well plates and exposed to increasing concentrations of cFLs (0 µM, 1 µM, 10 µM, 100 µM) for 30 minutes at 37 °C with continuous shaking. Subsequently, 20 µL of the 3-(4,5-dimethylthiazol-2-yl)-5-(3-carboxymethoxy-phenyl)-2-(4-sulfophenyl)-2H-tetrazolium (MTS) reagent (Abcam) was added to each well and incubated at 37 °C for 3 hours, according to the manufacturer's recommendation. The spectrophotometric absorbance of the samples was collected at 490 nm with a FLUOstar Omega plate reader (BMG Labtech). Data are presented as mean ± standard deviation and were calculated from a number of three repeats of three independent experiments.

#### Data analysis

### TIRF-M analysis of Fusion Events

Fusion events were manually detected in the raw video files as appearances of high-intensity signals in the liposome fluorescence channel that colocalized with a bacterium or the SLB plane. By selecting a 3x3 pixel area around the maximum intensity pixel (i.e. supposedly the attachment side), the normalised maximum fluorescence profiles over time have been extracted. The time-point of attachment was set to T=0 and the following 6 seconds were extracted for further analysis. For each fusion event, the following equation was fitted to the data:

$$I = exp(-\lambda t) + A$$

where I describes maximum fluorescence intensity and t is time, in order to extract the diffusion exponent λ and fluorescence plateau $A$. For further distinction between full-fusion and hemifusion events, we used a threshold of 0.5 for $A$ (hemifusion events being characterised as $A \geq 0.5$).

### Colocalization analysis

#### Mander's colocalization analysis between cytoplasmic DNA and Rh-PE

To quantify and visualise colocalization between the cFL and Gram-positive bacteria, we use an adjusted Mander's colocalization coefficient (MOC) of the cFL channel with the Gram-positive bacteria channel. First, both channels are independently otsu thresholded. As we are only interested in positions where we find localised cFL, we only filter for those positions in the Gram-negative bacteria channel (i.e. $\boldsymbol{B_i} = \boldsymbol{0}$ for $i$ with $cFL_i = 0$ and $B_i = B_i$ for $i$ with $cFL_i > 0$) where $\boldsymbol{B_i}$ denotes the intensity of the Gram-negative bacteria channel at pixel $i$ and $cFL_i$ the intensity of the cFL channel at pixel $i$, respectively. With this adjusted channel data, we then calculate the well established MOC as

$$MOC = \frac{\sum_i (B_i \times cFL_i)}{\sqrt{\sum_i B_i^2 \times \sum_i cFL_i^2}}.$$

### Spearman colocalization analysis between Vancomycin-BODIPY and Rh-PE

To quantify and visualise colocalization between the Vancomycin-BODIPY and Rh-PE, we use the Spearman rank correlation coefficient (SPCC). After otsu thresholding each channel, we then calculate the SPCC as

$$\boldsymbol{SPCC} = \mathbf{1} - \frac{\mathbf{6}\sum_i d_i^2}{\boldsymbol{n}(\boldsymbol{n}^2 - \mathbf{1})}$$

where $n$ is the number of pixels of the image and $d_i$ describes the difference in rank (ranked regarding pixel intensity) of the two channels for each pixel $i$.

### Degree of coverage analysis between Gram-negative Bacteria and cFLs

To analyze the extent of liposome coverage on Gram-negative BL21 cells, we determined the overlap between the bacterial outline and the liposomal Rh-PE signal. To achieve this, the reconstructed two-channel SIM images of the cytoplasmic GFP and liposomal Rh-PE were auto thresholded using Fiji [6]. Next, the outline of the cytoplasmic GFP signal was determined using the particle analyzer in Fiji on the thresholded GFP signal. The percentage of overlap between each outline and the thresholded Rh-PE signal was calculated using a custom-written Python code. The image processing pipeline is displayed in Supplementary Fig. 8.

### Modelled SIM data

For comparison of the experimentally acquired SIM data to a capsule model of the bacteria, artificial SIM images were generated from the approximate dimensions of the bacteria being imaged. To achieve this, SIM images of the cytoplasmic dye were first binarized and segmented to extract the length and width of the cross section. From these dimensions, an ideal stadium shape was created and rotated around the major axis to form the capsule model of the bacteria. The surface of this 3D volume was used as a model for the membrane and the interior of the volume was used as a model for the cytosol.

From the modelled 3D structures, artificial SIM images were generated by convolving the volume with a model point spread function (PSF) for the microscope used. This PSF was itself approximated from a basic model of the detection PSF and the SIM reconstruction parameters as reported from the SIM reconstruction software used.[7] Modelling the SIM process in this way is important as the reconstruction process imparts optical sectioning to the images which is not accounted for in a simplistic widefield model of the imaging system.[8] Line profiles across the minor axis of the resulting SIM images were then compared to the corresponding line profiles of the experimentally acquired SIM images. All image analysis and modelling was performed in MATLAB. The code underlying this work can be found on the GitHub repository: https://github.com/edward-n-ward/bacteria-modelling.

### Data visualization and statistical analysis

Graphs were plotted using Origin 2019. Statistical significance between two values was assessed using a two-tailed, unpaired Student's t-test (Origin 2019). Asterisks indicate statistical significance as determined by the Student's t-test (*$P < 0.05$, **$P < 0.01$, ***$P < 0.001$, and ****$P < 0.0001$).

**Video description**

**S1 Supplementary Video 1**

A representative TIRF-M video illustrates cFLs cFLs fusion with an *E.coli* supported lipid bilayer. An individual liposome arrives in the SLB plane. Rh-PE diffuses radially into the SLB. Scale bar: 5 µm.

**S2 Supplementary Video 2**

A representative TIRF-M video illustrates cFLs attachment to a BL21 bacterium. An individual liposome attaches to a bacterium without fusing. Scale bar: 1 µm.

**S3 Supplementary Video 3**

A representative TIRF-M video illustrates cFLs hemifusion with a BL21 bacterium. An individual liposome attaches to the bacterium. RH-PE diffuses into the bacterial membrane, leaving behind a high-intensity fluorescence at the attachment site. Scale bar: 1 µm.

**S4 Supplementary Video 4**

A representative TIRF-M video illustrates cFLs fusion with a BL21 bacterium. An individual liposome attaches to the bacterium. RH-PE diffuses into the bacterial membrane. Scale bar: 1 µm.

**Supplementary figures**

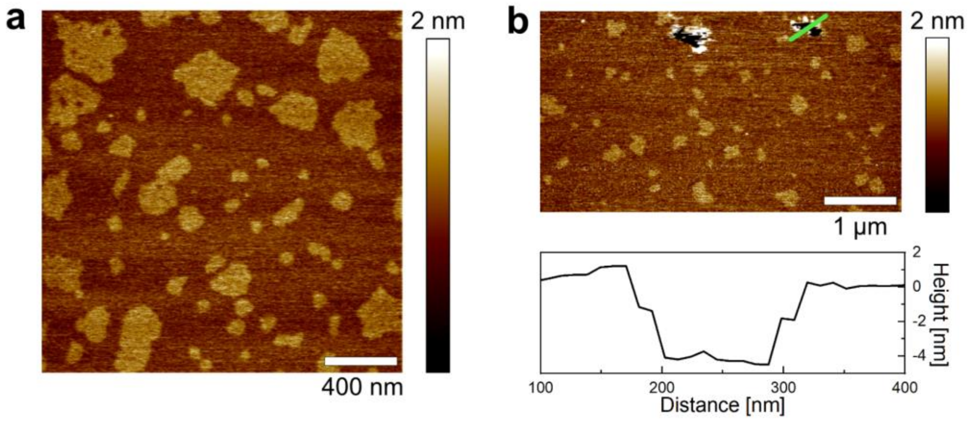


**Supplementary Figure 1:** *E.coli* total lipid extract supported lipid bilayers (SLB). (a) Representative atomic force microscopy (AFM) image of *E.coli* total lipid extract SLB, revealing lipid bilayer domains. (b) Representative AFM image of *E.coli* total lipid extract SLB with defects indicated by a red arrow. The bilayer has a surface height of approximately 4 nm which is in agreement with previously reported *E. coli* lipid bilayer heights.[9] The height profile corresponds to the green line in the AFM image.

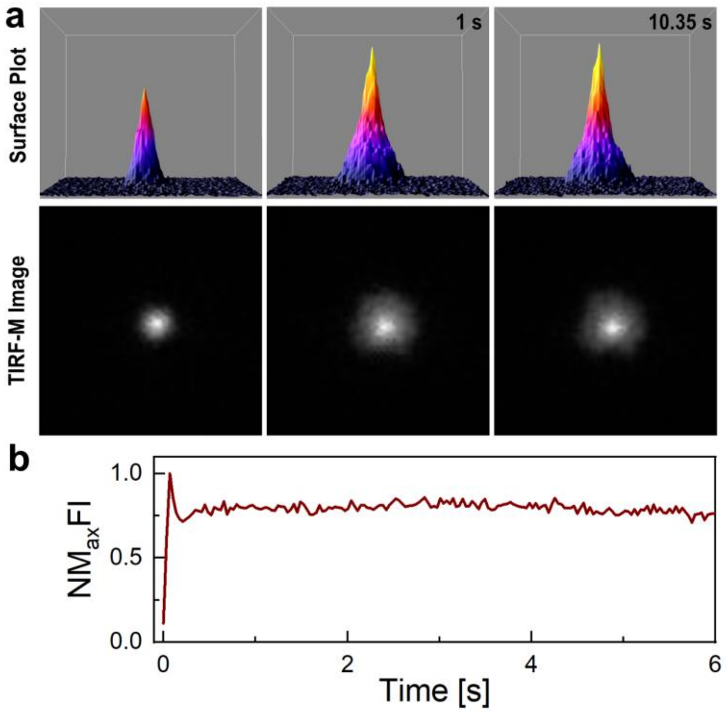


**Supplementary Figure 2:** Cationic fusogenic liposomes (cFL) attach to and spread on glass surfaces. (a) Time-lapse surface plot and corresponding total internal reflection microscopy (TIRF-M) images of cFLs spreading after encountering a glass substrate. (b) Normalised maximum fluorescence intensity (NMFI) profile plot of the cFLs in (a) within a 3x3 pixel area around the attachment point. The intensity profile shows an initial drop in the fluorescence intensity by approximately 25 % which is attributed to liposomal spreading on the glass surface. Negligible photobleaching is observed over a time period of 6 seconds.

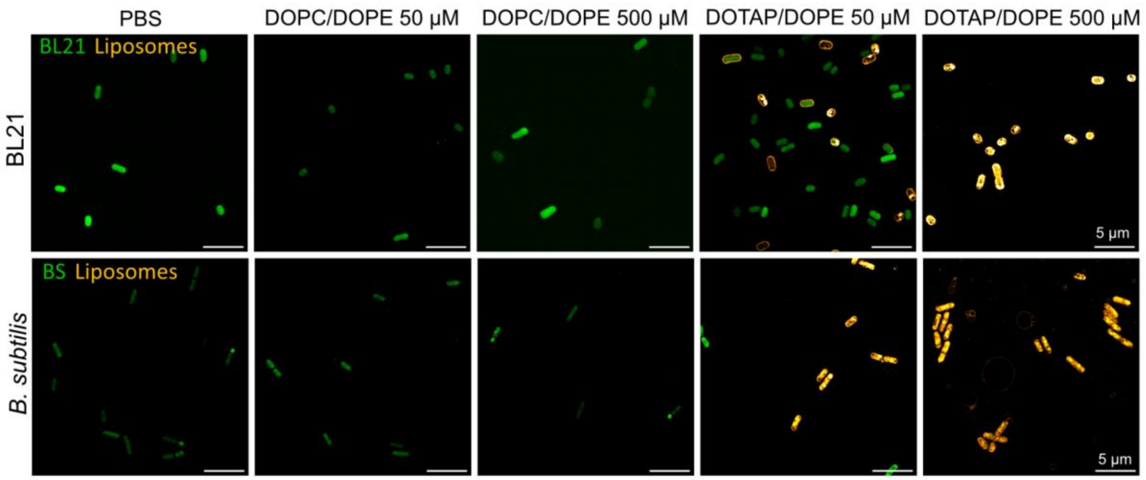


**Supplementary Figure 3:** DOTAP facilitates electrostatically driven bacterial targeting. Representative structured illumination microscopy (SIM) images of liposome exposed Gram-negative BL21 cells (top row) and Gram-positive *B. subtilis* cells (bottom row). The DOPC/DOPE (1:1 mole ratio) liposome composition in absence of the cationic lipid component DOTAP did not result in bacterial targeting.

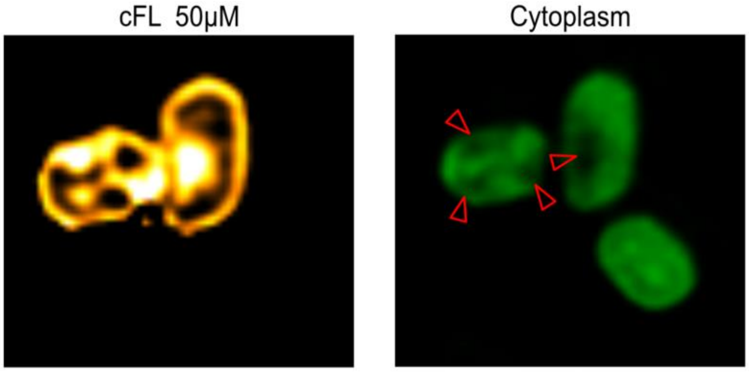


**Supplementary Figure 4:** Internal Rh-PE signal in BL21 cells is accompanied by GFP signal loss in the same area. Representative SIM images of cFLs targeted Gram-negative BL21 cells. Bacteria with internal Rh-PE signals (orange) show a loss in cytoplasmic GFP in the same area (highlighted by red arrows).

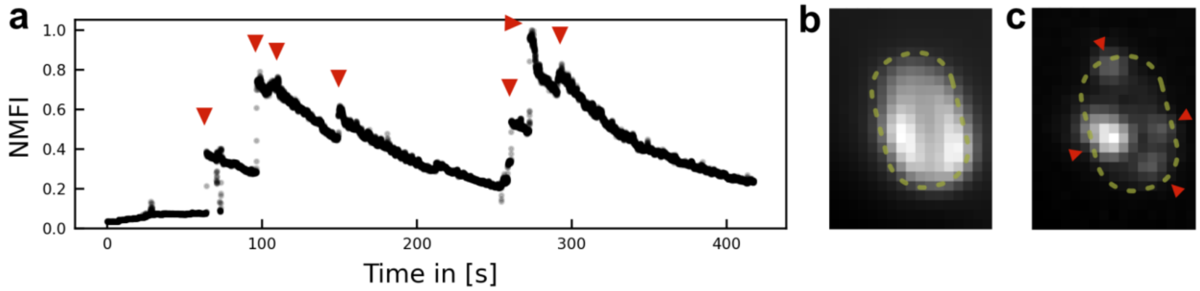


**Supplementary Figure 5:** Multiple liposomes attach to individual bacteria and integrate into their outer membrane. (a) A representative normalized maximum fluorescence intensity (NMFI) profile of the Rh-PE signal within the highlighted outline in (b), recorded over a period of approximately 6 minutes. Red arrows indicate liposome attachment events that increase the maximum recorded Rh-PE signal, followed by a decline in fluorescence intensity due to lipid dilution in the outer membrane and photobleaching effects. (b) A representative TIRF-M image of the Rh-PE signal of an E. coli cell upon liposome exposure. The yellow trace outlines the bacterium. (c) A representative Z-projection of the SIM slices that showed liposome attachment in (b). Red arrows indicate predominant liposome attachment sites.

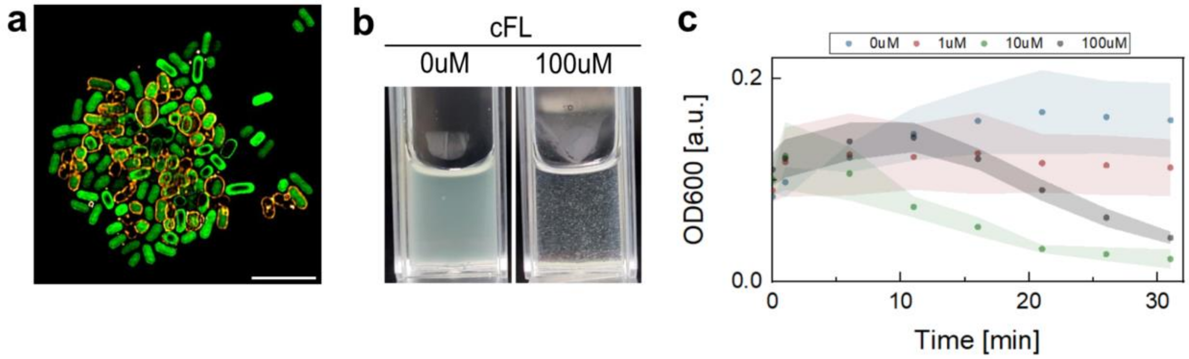


**Supplementary Figure 6:** Cationic lipid nanoparticles drive bacterial agglomeration. (a) Representative SIM image of BL21 agglomerates upon cFLs exposure. Scale bar, 5 µm. (b) Photograph of BL21 cell agglomeration in phosphate buffered saline (PBS) upon exposure to cFLs. (c) Optical density of bacterial cultures upon exposure to cFLs. The optical density of the suspension decays over time due to bacterial clumping. Presenting Mean ± SD based on 3 independent repetitions, each comprising three measurements.

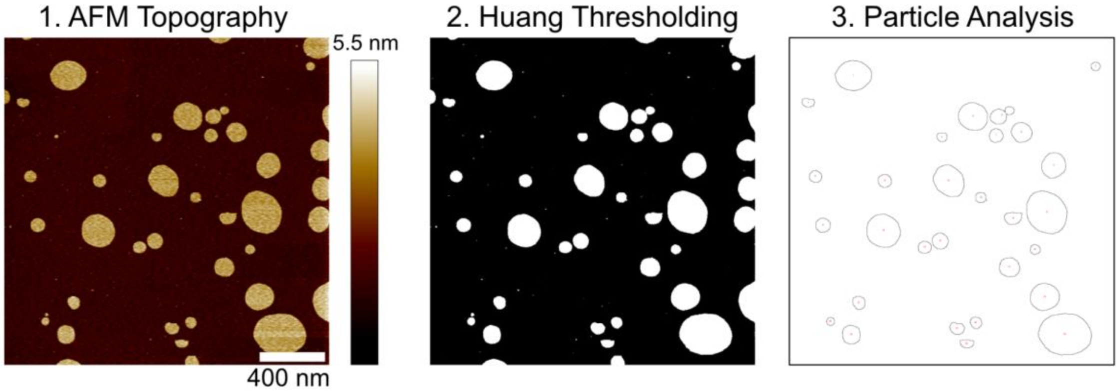


**Supplementary Figure 7:** Image processing pipeline to infer cFLs bilayer area on mica substrates. AFM topography images were saved in a tag image file format and processed using Fiji.[6] The images were thresholded using the Huang method [10] and the particle analyser was used to calculate the particle size. One pixel was measured to be 1.39 nm.

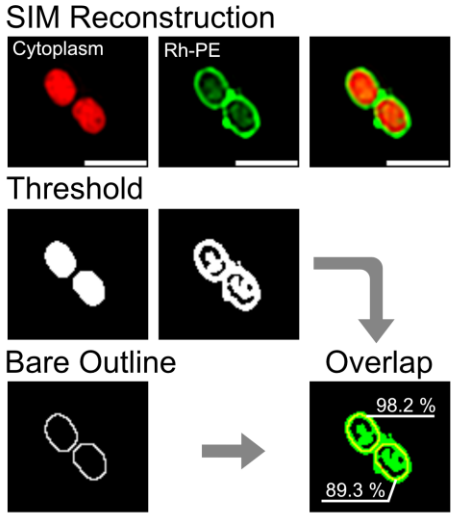


**Supplementary Figure 8:** Degree of coverage analysis between Gram-negative BL21 cell outline and cFLs. Reconstructed SIM images of cytoplasmic GFP and Rh-PE were thresholded using Fiji.[6] The outline of the cytoplasmic GFP signal was determined using the particle analyzer in Fiji and the overlap with the thresholded Rh-Pe signal was calculated.

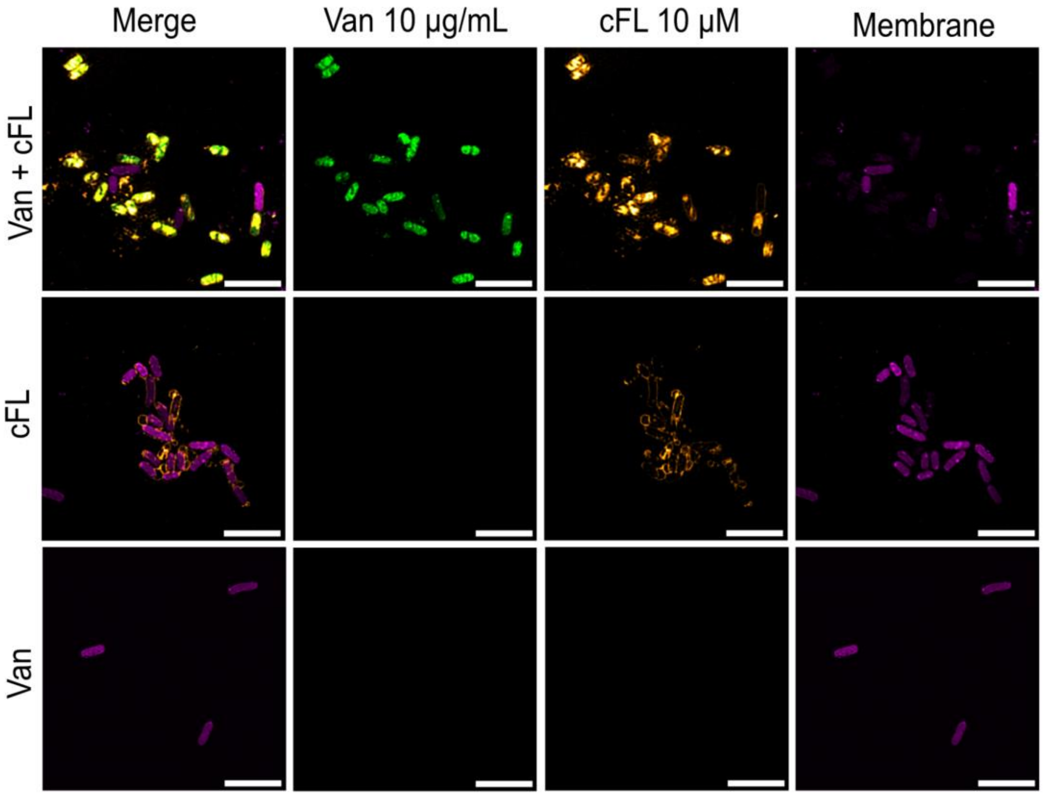


**Supplementary Figure 9:** Enhanced vancomycin targeting upon cFLs exposure. Representative SIM images of BL21 cells upon exposure to 10 μg/mL vancomycin (Van) co-delivered with 10 μM cFLs (top row), cFLs only (middle row), and vancomycin only (bottom row). Scale bar, 5 μm. BL21 cells were stained with 1 μM Mitotracker Far Red for membrane labeling.
